# Epidermal Stratification Requires Retromer-Mediated Desmoglein-1 Recycling

**DOI:** 10.1101/2021.10.25.464989

**Authors:** Marihan Hegazy, Jennifer L. Koetsier, Amber L. Huffine, Joshua A. Broussard, Brendan M. Godsel, Lisa M. Godsel, Kathleen J. Green

## Abstract

Sorting and trafficking transmembrane cargo is essential for tissue development and homeostasis. However, the importance of intracellular trafficking in the development and regeneration of stratified epidermis has not been investigated. Here we identify the interaction between VPS35, an essential component of the retromer endosomal trafficking complex, and the desmosomal cadherin, Desmoglein 1 (Dsg1). Dsg1 is specifically expressed in stratified tissues and when properly localized on the plasma membrane, promotes epidermal stratification. We show that the retromer drives Dsg1 recycling from the endo-lysosomal system to the plasma membrane to support keratinocyte stratification and differentiation. The retromer enhancing chaperone, R55 promotes the plasma membrane localization of Dsg1 and a Dsg1 mutant associated with Severe dermatitis, multiple Allergies, and Metabolic wasting (SAM) syndrome, enhancing the ability of SAM-Dsg1 to promote stratification. Our work provides the first evidence for retromer function in epidermal regeneration and identifies it as a potential therapeutic skin disease target.

## INTRODUCTION

The epidermis is a complex stratified tissue composed predominantly of keratinocytes that undergo a highly orchestrated process of differentiation and stratification to create an essential barrier that protects against water loss, pathogens, and allergens (Proksch et al., 2008). The formation and maintenance of the epidermal barrier is dependent on intercellular junctions, including gap junctions, tight junctions, adherens junctions (AJs), and desmosomes (Garcia et al., 2018; Rubsam et al., 2018; Sumigray and Lechler, 2015). AJs and desmosomes are structurally similar with transmembrane adhesive proteins called cadherins that associate with armadillo and cytoskeletal adaptor proteins to anchor actin or intermediate filaments to the plasma membrane, respectively (Green et al., 2020; Saito et al., 2012). Desmosomes are required to maintain epidermal integrity and modulate cell mechanics in tissues subjected to high mechanical stress such as the skin and heart (Garrod and Chidgey, 2008; Hatzfeld et al., 2017; Hegazy et al., 2021; Zimmer and Kowalczyk, 2020).

Desmosome composition is different in each layer of the epidermis, and this patterning is essential for the functional polarity of the multi-layered tissue. Among the most highly patterned molecules in the epidermis are the intercellular adhesion proteins, the desmosomal cadherins. The desmosomal cadherins are divided into two subclasses, desmogleins and desmocollins, for a total of at least seven protein isoforms. One such isoform, desmoglein-1 (Dsg1), is first expressed as keratinocytes commit to differentiate and transit into the next superficial layer on their journey to the skin’s surface. Its expression gradually increases throughout differentiation, reaching its highest level in the outer living layers of the epidermis (Rubsam et al., 2018). Dsg1 distribution is critical, in turn, for patterning ErbB family signaling pathways in the epidermis; it inhibits EGFR/MAPK signaling during early differentiation and promotes ErbB2-mediated tight junction barrier formation during late differentiation (Broussard et al., 2021; Getsios et al., 2009; Harmon et al., 2013). Dsg1’s in vivo importance is reflected in the observed disruption of signaling pathways controlling differentiation, barrier and immune function in Dsg1-deficient animal models (Godsel et al., 2020; Kugelmann et al., 2019). In addition, Dsg1 deficiencies result in human inherited and autoimmune diseases including Severe dermatitis, multiple Allergies, and Metabolic wasting (SAM) syndrome, striate palmoplantar keratoderma (SPPK), and pemphigus foliaceus (PF) (Cheng et al., 2016; Dua-Awereh et al., 2009; Has et al., 2015; Hershkovitz et al., 2009; Samuelov et al., 2013; Schmidt et al., 2019).

In addition to its role in patterning ErbB family signaling, we recently demonstrated that Dsg1 promotes stratification during the basal to suprabasal commitment to differentiate. It does so by remodeling cortical actin to redistribute membrane tension in basal cells (Nekrasova et al., 2018). This redistribution is required for releasing basal cells from the basement membrane for transit into the suprabasal layers, a process called delamination that contributes to the formation and maintenance of the regenerating multi-layered epidermis (Damen et al., 2021; Miroshnikova et al., 2018; Nekrasova et al., 2018; Watt and Green, 1982; Williams et al., 2014). Dsg1’s role in this process requires its proper localization on the plasma membrane, in detergent insoluble regions segregated from the AJ protein, E-cadherin (Nekrasova et al., 2018). Inhibiting Dsg1’s association with the dynein light chain, Tctex-1, disrupts its proper membrane localization, actin remodeling and thus, basal cell delamination. Further, mutations that impair accumulation of Dsg1 on the plasma membrane have been shown to cause the severe systemic inflammatory disorder, SAM syndrome (Lewis et al., 2019; Samuelov et al., 2013). While these observations illustrate the importance of Dsg1 localization for normal epidermal homeostasis, there are substantial gaps in our understanding of the mechanisms by which Dsg1 is delivered to its final destination on the plasma membrane.

To identify trafficking machinery that regulates Dsg1 plasma membrane localization, we performed two independent BioID protein interaction screens and discovered VPS35, a component of a trafficking complex called the retromer, as a potential Dsg1 interacting protein. The retromer functions in endosomal trafficking, including retrograde trafficking from the endosome-to-*trans*-Golgi network (TGN), polarized transport of membrane proteins, and endosome-to-plasma membrane recycling (Seaman, 2012). It is composed of a trimer consisting of VPS35-VPS26-VPS29 with a role in cargo selection, and a Snx component, which plays a role in promoting the formation of dynamic tubulovesicular structures for protein transport (Burd and Cullen, 2014). While a role for retromer dysfunction in neuronal diseases such as Parkinson’s, Alzheimer’s and amyotrophic lateral sclerosis (ALS) disease is emerging (Carosi et al., 2021), information about retromer in the epidermis is limited to a role in Human Papilloma Virus (HPV) trafficking (Lipovsky et al., 2013). Here we demonstrate that the retromer promotes Dsg1 recycling and accumulation on the plasma membrane to promote epidermal stratification, which is enhanced by a small molecule retromer chaperone called R55. In addition, R55 restores the plasma membrane localization of SAM syndrome-associated Dsg1 with a mutation in its signal sequence, increasing its ability to promote stratification to the level of WT-Dsg1. These data raise the possibility that retromer chaperones may be used clinically in the future to help ameliorate symptoms in Dsg1 deficiency disorders.

## RESULTS

### Inhibition of endosomal trafficking disrupts plasma membrane localization of Dsg1, but not the AJ cadherin, E-cadherin

Dsg1 incorporates into desmosomes that are segregated from E-cadherin in AJs to promote actin cytoskeletal reorganization in cells committing to differentiate (Nekrasova et al., 2018). This segregation could occur at least in part through utilization of distinct trafficking mechanisms by Dsg1 and E-cadherin. To identify potential mediators of differential trafficking, we carried out parallel BioID screens using Dsg1 and E-cadherin biotin ligase BirA* fusion proteins expressed in differentiated keratinocytes (Roux et al., 2012). The retromer component, VPS35 was identified in the Dsg1, but not the E-cadherin, screen. VPS35 was also present in a second, independently performed screen. Identification of VPS35 as a potential binding partner was the first indication that Dsg1 may require endosomal-mediated transport for its plasma membrane localization. To test the importance of endosomal trafficking on Dsg1 plasma membrane localization, we treated differentiated keratinocytes with primaquine, a weak base that prevents proteins from exiting the endosome (van Weert et al., 2000). Addition of primaquine resulted in the cytoplasmic accumulation of endogenous and ectopically expressed Dsg1 in early endosomes identified by staining with EEA1 (Figure 1A, B). In addition, Dsg1 levels at the cell-cell interface were significantly diminished from vehicle control (Fig. 1A, C and Figure S1C, D, H). 3D reconstruction of confocal images illustrates Dsg1 impairment in the stratified, upper layers where Dsg1 is concentrated (Figure 1A, D, H and Figure S1E). Primaquine does not affect endogenous and ectopically expressed E-cadherin localization or colocalization with EEA1, whether it is added after the initiation of differentiation or when introduced to undifferentiated cells (Figure 1E-H and Figure S1A-D, F-H). Thus, inhibition of endosomal trafficking with primaquine disrupts Dsg1, but not E-cadherin, localization resulting in its accumulation in endosomes. These data suggest that Dsg1’s accumulation in the desmosomes of keratinocytes requires endosomal-mediated trafficking steps not used by E-cadherin.

**Figure 1.**
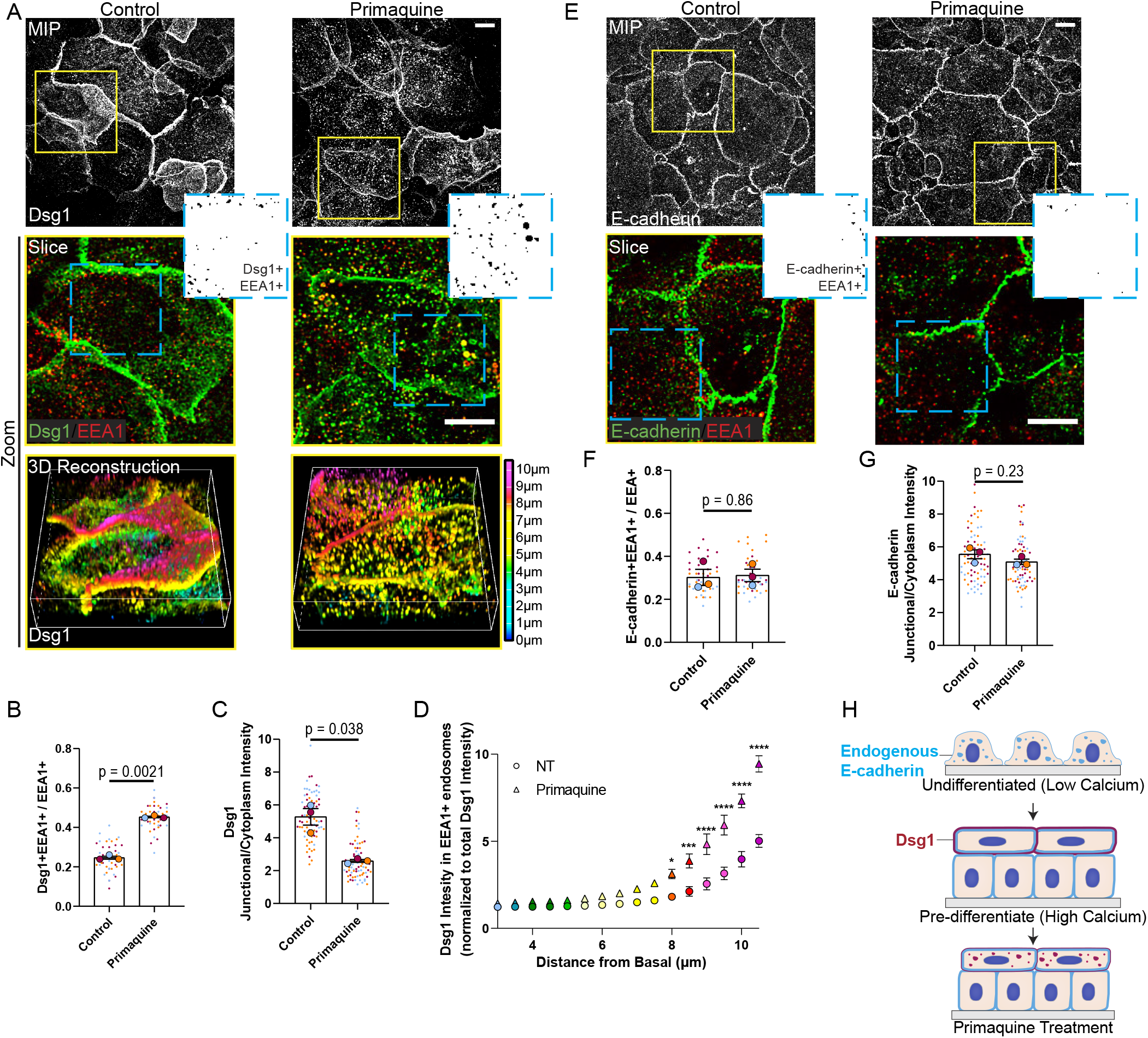
Dsg1 requires endosomal trafficking for its plasma membrane localization. (A, E) Top (MIP): Maximum intensity projection (MIP) of Dsg1 (A) and E-cadherin (E) immunofluorescence following overnight Primaquine treatment (200µM) of pre-differentiated keratinocytes. Scale bar, 20 µm. Middle (Slice): 1 µm slice of confocal images with dual staining of EEA1 pseudocolored red with Dsg1 (A) or E-cadherin (E) in green. Scale bar, 20 µm. Inset: Auto-threshold was implemented using ImageJ for EEA1 and Dsg1 (A) or E-cadherin (E). Black puncta illustrate the region of overlap between EEA1 signal with Dsg1 (A) or E-cadherin (E) signal. Bottom of A (3D reconstruction): 3D reconstruction of Dsg1 with a z-depth pseudocoloring illustrating Dsg1 localization in the stratified layer (See Supplemental Figure 1E for corresponding DAPI co-stain for Dsg1and Supplemental Figure 1F for 3D reconstruction of E-cadherin). (B, F) Fraction of EEA1+ vesicles that are Dsg1+ (B) or E-cadherin+ (F) per field; paired t-test from three biological repeats, and error bars are SEM. (C, G) Quantification of average Dsg1 (C) and E-cadherin (G) membrane/cytoplasmic ratio following Primaquine treatment; paired t-test from three biological repeats, and error bars are SEM. (D) Quantification of Dsg1 intensity in EEA1 positive vesicles with respect to the distance from the basal layer (colors correspond to z-depth look up table in (A)); two-way ANOVA with Sidak post-hoc test from three biological repeats, and error bars are SEM (See Supplemental Figure 1G for E-cadherin equivalent). (H) Summary of the impact of Primaquine treatment on endogenous Dsg1 (colored red on the plasma membrane in stratified cells and in vesicles in Primaquine treated cells and) and E-cadherin (colored blue in vesicles and the plasma membrane) localization after the initiation of keratinocyte differentiation.

### Dsg1 interacts with the essential retromer component VPS35

As mentioned above, VPS35 was detected in two BioID protein interaction screens using Dsg1-BirA*. The canonical function for VPS35 in the retromer is cargo recognition. To test if Dsg1 can be recognized as cargo by the retromer, we immunoprecipitated VPS35 in keratinocytes expressing Dsg1-FLAG and evaluated proteins in the complex by immunoblotting. Dsg1-FLAG co-immunoprecipitated with VPS35, whereas E-cadherin was only weakly detectable (Figure 2A). Cation-independent Mannose-6 Phosphate Receptor (CI-M6PR), which is known to undergo retromer-dependent trafficking from the endosome-to-TGN, was used as a positive control and co-immunoprecipitated with VPS35 as expected (Arighi et al., 2004).

**Figure 2.**
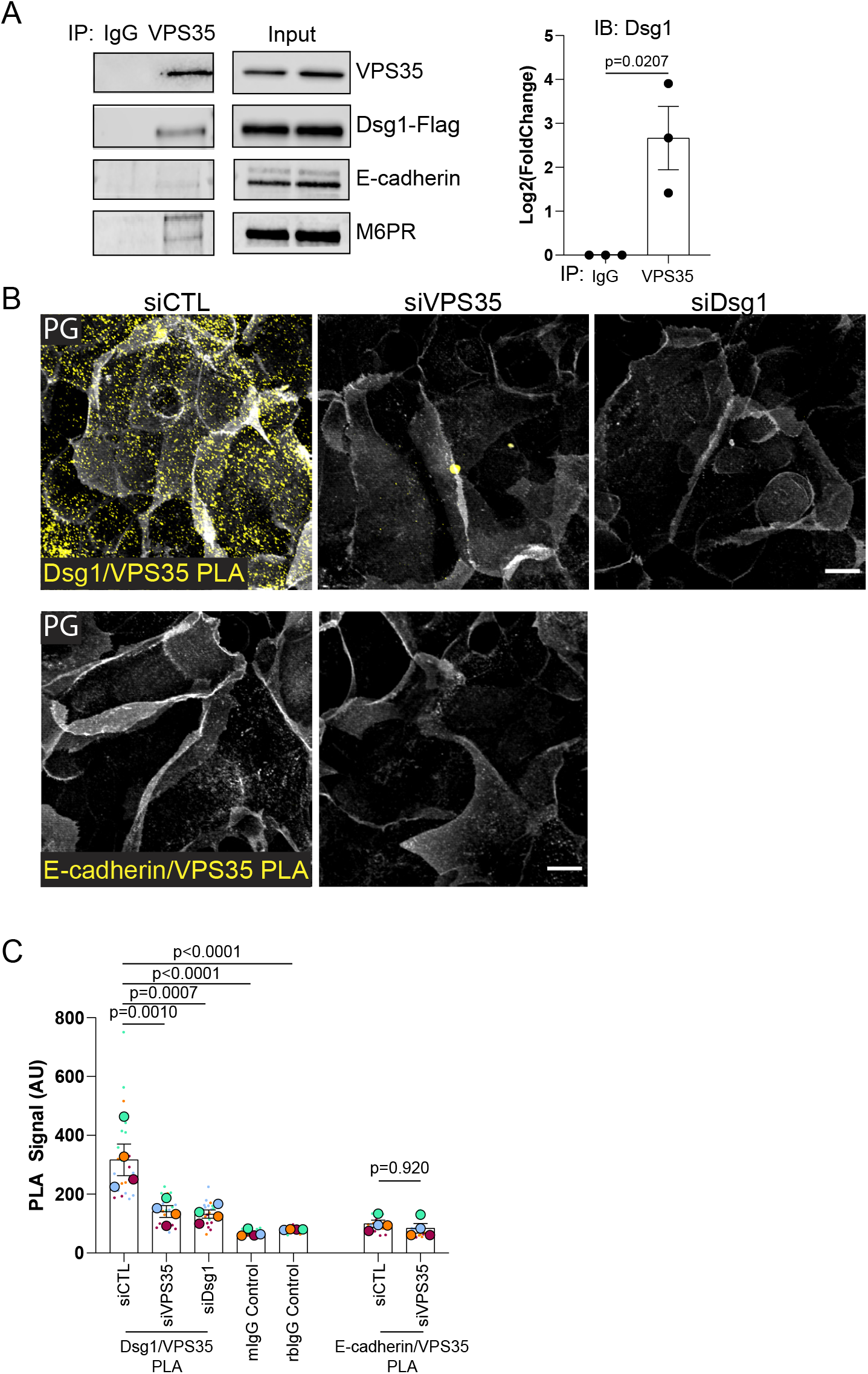
The essential retromer component VPS35 interacts with Dsg1. (A) Co-immunoprecipitation using an anti-VPS35 antibody and immunoblotting for Dsg1-FLAG, E-cadherin, and M6PR (positive control). Quantification of the densitometry of the Dsg1 immunoblot on the right; unpaired two-way t-test from three biological repeats, and error bars are SEM. (B) Proximity Ligation Assay (PLA) using an anti-VPS35 antibody paired with an anti-Dsg1 or anti-E-cadherin antibody to test the proximity of VPS35 with endogenous Dsg1 or E-cadherin in situ. siRNA-mediated depletion of Dsg1 and/or VPS35 was used to test for specificity of the PLA signal and compared to scramble control (siCTL). See Figure S2 for additional controls. Plakoglobin (PG) staining to visualize cell-cell membranes (white). Scale bar, 20 µm (C) Quantification of the Dsg1-VPS35 PLA and E-cadherin-VPS35 PLA compared to IgG control and/or knockdown control; one-way ANOVA with Dunnett post-hoc test (Dsg1-VPS35 PLA) or paired t-test (E-cadherin-VPS35 PLA) from four biological repeats, and error bars are SEM.

To verify that endogenous Dsg1 and VPS35 are in close proximity in situ, a Proximity Ligation Assay (PLA) was performed on differentiated keratinocytes. A Dsg1/VPS35 antibody pairing produced a positive PLA signal in stratified cells where Dsg1 is highly concentrated, when compared with IgG or siDsg1 or VPS35 RNAi-treated controls (Figure 2B, C and Figure S2A). An E-cadherin/VPS35, antibody pairing did not produce a PLA signal in siControl or siVPS35-treated cells, in contrast to an E-cadherin/β-catenin antibody pairing, which produced a strong PLA signal (Figure 2B, C and Figure S2A). Tandem immunofluorescence staining and immunoblotting confirmed antibody recognition and knockdown efficiency in each experiment (Figure S2B-D). These data suggest there is a pool of Dsg1 in close proximity to the retromer component VPS35 in stratified keratinocytes.

### Loss of the retromer results in the accumulation of Dsg1 in the endo-lysosomal system

The data showing that Dsg1 is found in a complex with VPS35 (Figure 2), and that endosomal trafficking is necessary for Dsg1 plasma membrane localization (Figure 1) suggest that Dsg1 may utilize the retromer for trafficking to the plasma membrane. To test this hypothesis, VPS35 and another essential retromer component, VPS29, were depleted in cells and the amount of plasma membrane localized Dsg1 was quantified. Loss of either retromer component resulted in a decrease in surface localized Dsg1, while no change to E-cadherin surface levels was observed (Figure 3A, B and Figure S3A). The retromer is necessary for the tubulation and fission of endosomes to promote cargo exit from the lysosomal degradative pathway (Burd and Cullen, 2014; van Weering et al., 2012). Live cell imaging of keratinocytes revealed highly dynamic Dsg1-GFP positive tubules extending from vesicles, which became more rounded and less elongated in cells silenced for either VPS35 or VPS29 (Figure 3C, D and Movie 1). These data suggest that the retromer is necessary for Dsg1 localization in endosomal tubules and exit out of the endosome.

**Figure 3.**
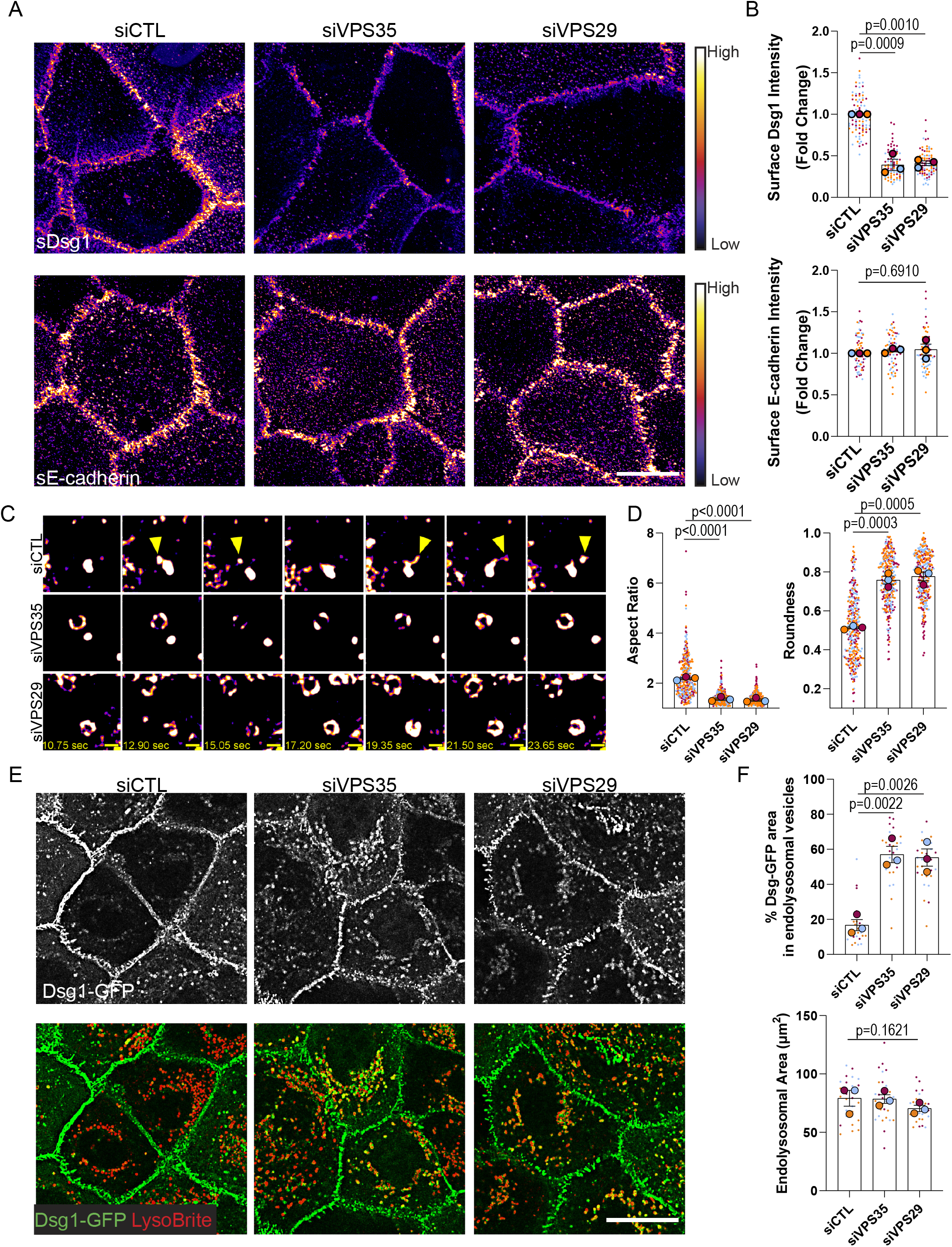
Depletion of retromer components disrupts Dsg1 plasma membrane localization, endocytic vesicle tubulation, and results in Dsg1 accumulation in the endo-lysosomal system. (A) Living cells were surface-labeled for Dsg1 or E-cadherin following siRNA-mediated depletion of VPS35 or VPS29 and compared to scramble transfected control (siCTL). Images were pseudocolored using ImageJ with a Fire look up table. Scale bar, 20 µm. (B) Quantification of junctional Dsg1 or E-cadherin; one-way ANOVA with Dunnett post-hoc test from three biological repeats, and error bars are SEM. (C). Movie stills from Dsg1-GFP live cell imaging with yellow arrowheads highlighting Dsg1 localization in vesicle tubules and projections. Pseudocol ored as in (A). Refer to Supplemental materials to view the movie. Scale bar, 1 µm. (D) Quantification of aspect ratio and roundness of Dsg1-GFP vesicles; one-way ANOVA with Dunnett post-hoc test from three biological repeats, and error bars are SEM. (E) Immunofluorescence of Dsg1 and endo-lysosomal marker, LysoBrite, in live cells following retromer component depletion compared to scramble control (siCTL). Scale bar, 20 µm. (F) Quantification of the % area of Dsg1 in endo-lysosomal vesicles and the area of LysoBrite labeled endo-lysosomes; one-way ANOVA with Dunnett post-hoc test from three biological repeats, and error bars are SEM.

To test if Dsg1 becomes trapped in the endo-lysosomal system upon loss of the retromer, LysoBrite was used to label acidic organelles such as the endosome and lysosome. Consistent with the idea that Dsg1 requires the retromer for its exit from the endosome, depletion of the retromer components, VPS35 or VPS29, resulted in the accumulation of ectopic Dsg1-GFP in the endo-lysosomal system in basal keratinocytes (Figure 3E, F). Similarly, in stratified cells, endogenous Dsg1 displayed enhanced colocalization with the endo-lysosomal marker, LAMP1, following retromer component depletion (Figure S3B). As a positive control, Glucose transporter 1 (GLUT1), another protein that utilizes the retromer for its endosome-to-plasma membrane trafficking, exhibited increased localization in LAMP1-positive vesicles in the basal layer of primary keratinocytes (Figure S3B) (Steinberg et al., 2013).

Efficient knockdown of retromer components was verified using immunoblotting (Figure S3A, C). Similar to what has previously been reported, expression of VPS35 or VPS29 is necessary to maintain protein levels of the other retromer components (Fuse et al., 2015). While it might be predicted that Dsg1 levels would decrease due to sequestration and degradation in lysosomes, we saw such a decrease inconsistently. The fact that loss of the retromer has been reported to reduce lysosomal proteolytic capacity could explain why total levels of Dsg1 did not consistently decrease despite being trapped in the endo-lysosomal system (Cui et al., 2019). Interestingly, retromer depletion in Dsg1-GFP expressing keratinocytes occasionally resulted in increased extracellular cleavage, which could reflect lysosomal or MMP/ADAM dependent cleavage (Figure S3C). Altogether, these data suggest that Dsg1 requires the retromer for its exit from the endo-lysosomal system and for its plasma membrane localization.

### Dsg1 utilizes the retromer for endosome-to-plasma membrane recycling

The retromer has been reported to promote endosome-to-TGN retrograde trafficking and endosome-to-plasma membrane trafficking connected with the recycling of some retromer cargo (Seaman, 2012). Based on the observation that loss of the retromer resulted in decreased levels of Dsg1 at the plasma membrane, we tested the role of the retromer in recycling the desmosomal cadherin to the plasma membrane using an antibody-based recycling assay (Figure 4A). After surface labeling cells with an anti-Dsg1 monoclonal antibody (I), internalization was induced using low calcium media with the calcium chelator EGTA for 30 minutes at 37°C (II). Following internalization, remaining surface molecules were blocked using an unconjugated goat anti-mouse F(ab)_2_ (III). The cells were then returned to 37°C in high calcium media to induce recycling, and this pool was recognized by surface staining with a 488-Alexa Fluor conjugated goat anti-mouse secondary antibody (IV). Internalized Dsg1 was visualized with a 568-Alexa Fluor conjugated goat anti-mouse secondary antibody after fixation and permeabilization (IV). Similar to Figure 3A, surface Dsg1 was reduced in retromer depleted cells at steady state, resulting in a 2-fold higher pool of labeled Dsg1 in control compared with VPS35 or VPS29 knockdown cells (Figure 4B, C). Correspondingly, an increased level of Dsg1 was observed in the internalized pool in retromer depleted cells when normalized to the starting surface pool. This increase might be explained by retention of Dsg1 in vesicles in the absence of the retromer instead of freely cycling back to the plasma membrane (Figure 4B, D). Finally, the plasma membrane associated recycled pool also decreased significantly upon loss of the retromer (Figure 4B, E, F). These data are consistent with the hypothesis that the retromer is necessary for Dsg1 endosomal recycling to the plasma membrane.

**Figure 4.**
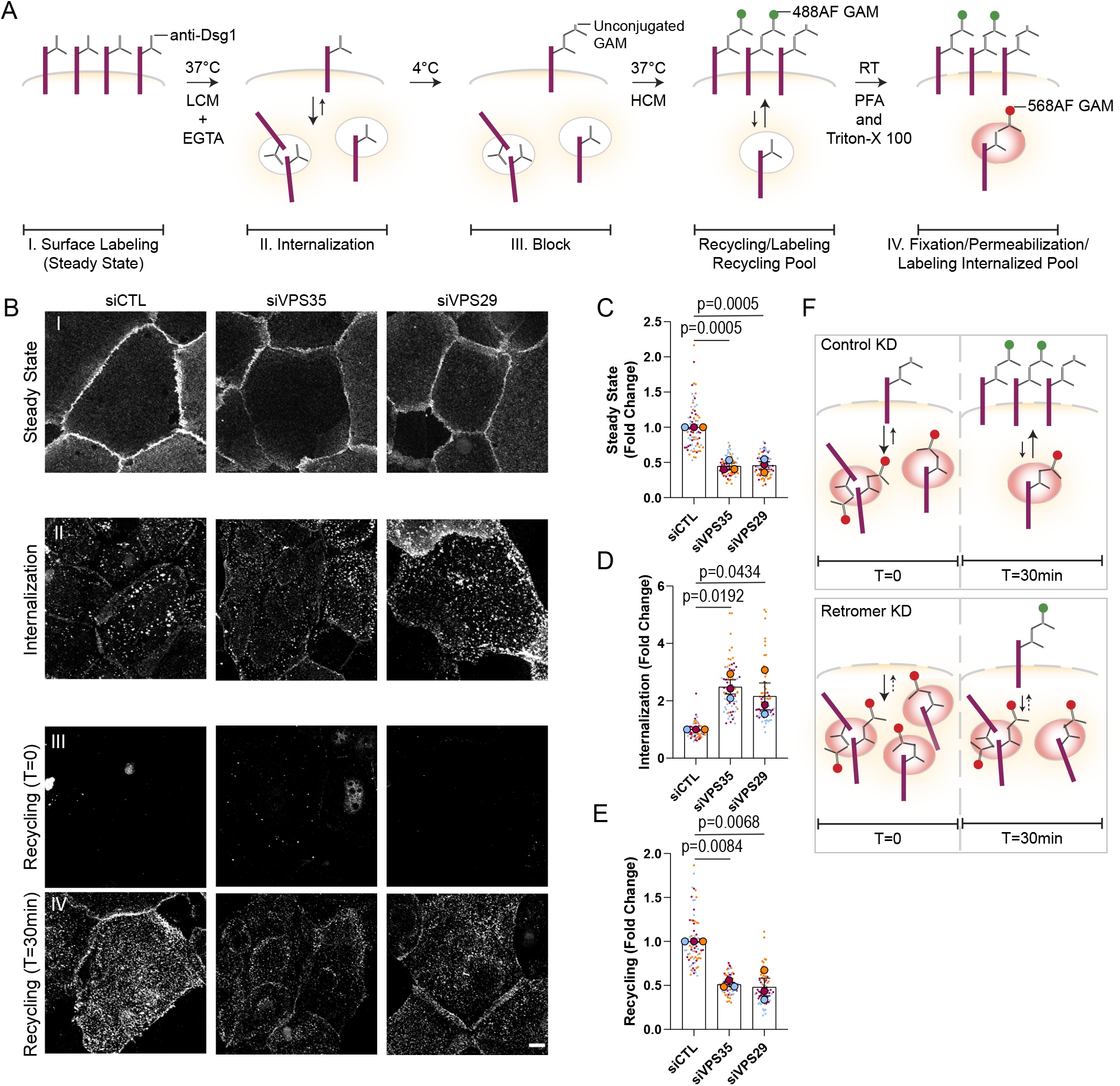
The retromer is necessary for Dsg1 endosomal-to-plasma membrane recycling. (A) Schematic of the antibody-based recycling assay shown in B. (A, B) I. Dsg1 surface signal at steady state with or without siRNA-mediated depletion of retromer components. II. Dsg1 internalization signal in the 568 channel following fixation and permeabilization after 30min low calcium and EGTA treatment. III. Dsg1 remaining surface signal in the 488 channel after 30min low calcium and EGTA treatment (T=0). IV. Dsg1 recycling signal after 30min high calcium treatment following internalization. Scale bar, 10 µm. (C) Quantification of Dsg1 surface signal at steady state (D) Quantification of Dsg1 internalization signal. (E) Quantification of Dsg1 recycling signal (C-E) Statistics performed with one-way ANOVA with Dunnett post-hoc test from three biological repeats, and error bars are SEM (F) Summary of the antibody-based recycling assay results showing that retromer loss impairs Dsg1 surface signal at steady state and enhances Dsg1 internalization due to the lack of constitutive recycling and disrupts Dsg1 recycling.

### Loss of the retromer results in abnormal differentiation

While Dsg1’s function in epidermal differentiation has been established, the retromer’s role in this process has not been addressed. To test if depletion of the retromer affects epidermal differentiation, the human epidermis was reconstituted using a 3D organotypic epidermal “raft” culture model where primary keratinocytes are lifted to an air-liquid interface on a collagen/fibroblast plug (Figure S4A)(Arnette et al., 2016). In control rafts, the basal, spinous, granular, and cornified layers are easily identified and resemble human skin epidermis 6-7 days after lifting the primary keratinocytes. When retromer components were depleted using siRNA, H&E staining revealed a disruption in structural features of the granular and cornified layers, including a loss of keratohyalin granules and loss of the superficial cornified layer (Figure 5A). We next stained raft cultures at 3 and 6-7 days after lifting to an air liquid interface for Dsg1 and Keratin 10 (K10), an intermediate filament protein expressed only in suprabasal cells. In day 3 rafts, the area of coverage of Dsg1 and K10 was reduced by half and staining was frequently absent from the most superficial layers where Dsg1 and K10 are typically expressed. Based on our previous observation that Dsg1 promotes K10 expression (Getsios et al., 2009), areas occupied by K10 also stained for Dsg1 (Figure 5B, C). The appearance of basal-like cells in the superficial layers was also observed, impairing the regular stacking of the multi-layered tissue and suggesting a disruption in tissue polarity (Figure 5B). Day 6-7 rafts exhibited a partial rescue in Dsg1 and less so K10 expression, as VPS35 expression began to return following knockdown (Figure 5D, E and Figure S4B). These data suggest that loss of the retromer impairs epidermal differentiation.

**Figure 5.**
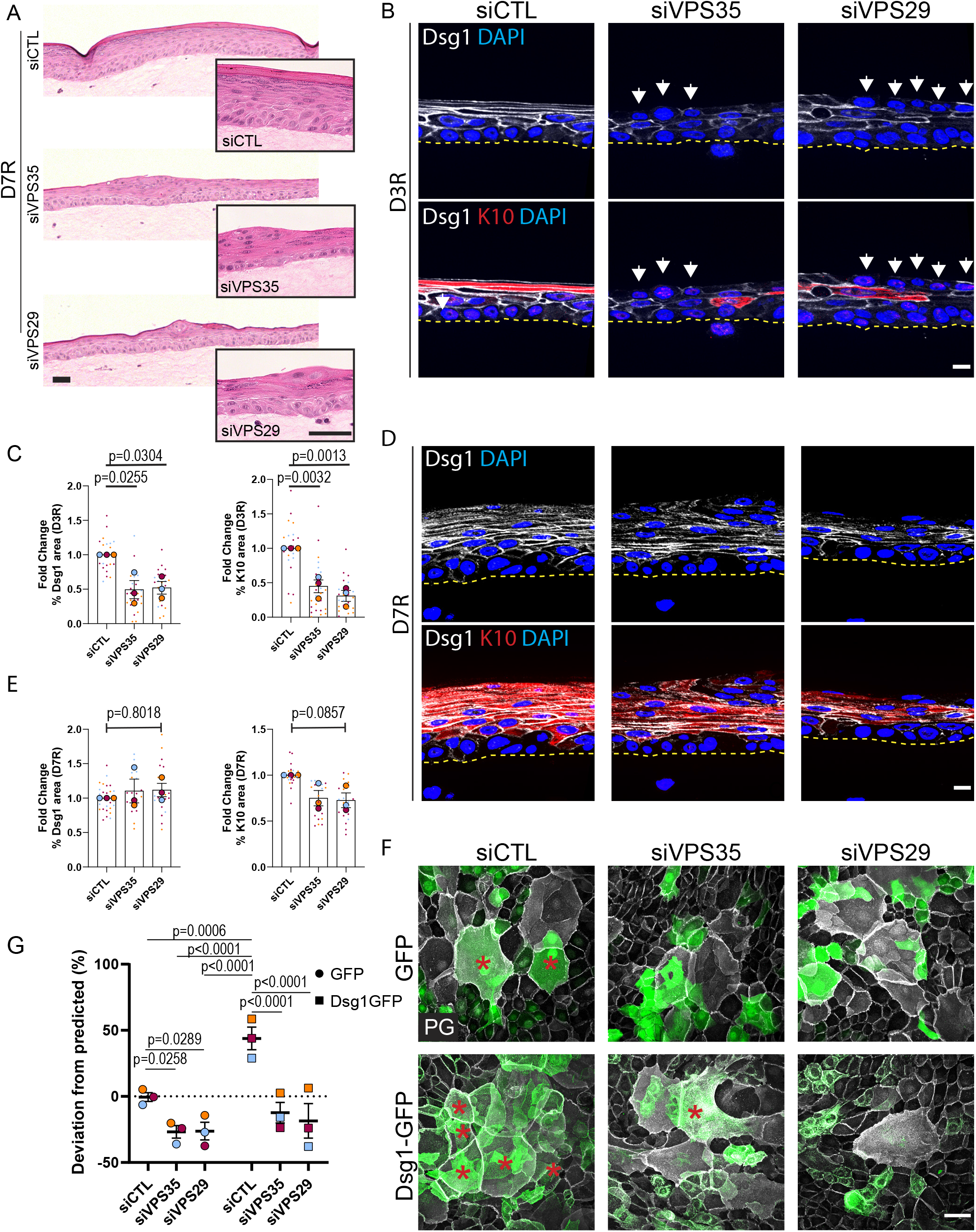
Retromer depletion disrupts epidermal differentiation, morphogenesis, and Dsg1-mediated stratification. (A) H&E staining of 3D organotypic “raft” cultures that were harvested 7 days after lifting to an air-liquid interface (D7R). Scale bar, 100 µm. Inset: Representative higher magnification H&E images. Scale bar, 50 µm. (B, D) Tissue sections from 3D rafts stratified for 3 days (D3R) (B) or 7 days (D7R) (D) were stained for Dsg1 and the differentiation marker Keratin 10 (K10). White arrows highlight basal-like cells in the superficial layers. Scale bar, 10 µm. (C, E) Quantification of % Dsg1 area and % K10 area per field in D3R (C) and D7R (E); one-way ANOVA with Dunnett post-hoc test from three biological repeats, and error bars are SEM. (F) GFP- and Dsg1-GFP labeled keratinocytes were transfected with siCTL, siVPS35, or siVPS29 before being mixed 50:50 with unlabeled keratinocytes and allowed to settle overnight. Cells were switched to 1.2mM CaCl_2_ media for 24hrs then fixed and stained with Plakoglobin (PG)/DAPI. Scale bar, 50 µm. (G) Quantification of deviation from predicted % of GFP cells or Dsg1-GFP cells in the stratified layer; one-way ANOVA with Tukey post-hoc test from three biological repeats, and error bars are SEM.

As Dsg1 promotes delamination of basal cells into the next superficial layer, we hypothesized that the observed impairment of differentiation may be due in part to a reduction in stratification due to impaired trafficking of Dsg1 in retromer-deficient cells (Nekrasova et al., 2018). To test whether retromer is required for efficient Dsg1-mediated stratification, we carried out a sorting assay that measures the ability of genetically modified GFP labeled cells to stratify when mixed in a 50:50 ratio with unlabeled keratinocytes (Broussard et al., 2021). Depletion of retromer components in GFP positive cells resulted in a decrease in the number of GFP positive keratinocytes that moved into the superficial layer, compared to the controls. This could be due to loss of endogenous Dsg1 from the plasma membrane or due to a disruption of another retromer cargo component important for stratification (Figure 5F, G). Therefore, to further test if the retromer is necessary for Dsg1-mediated stratification, we ectopically expressed Dsg1-GFP in basal keratinocytes, which we previously showed promotes differentiation and stratification (Broussard et al., 2021). As expected, Dsg1-GFP enhanced stratification; however, siRNA-mediated depletion of the retromer components blocked this increase (Figure 5F, G). These data support the hypothesis that the retromer is necessary for Dsg1-mediated keratinocyte stratification.

### The retromer chaperone R55 promotes VPS35 association with and plasma membrane localization of wild type (WT) and disease associated Dsg1

Bi-allelic loss of function mutations in Dsg1 are associated with a severe systemic inflammatory disorder known as SAM syndrome. One of the originally reported mutations, c.49-1G>A, causes skipping of exon 2 in a region with 0.981 probability of encoding a signal peptide, which is important for targeting transmembrane molecules to the endoplasmic reticulum (ER) (Figure S5A, B) (Almagro Armenteros et al., 2019). The c.49-1G>A SAM mutant has a 0.5477 probability of having a signal peptide (Figure S5A, B). This SAM mutant is expected to inefficiently insert into the ER membrane, thus decreasing the pool of mature Dsg1 available to traffic to the plasma membrane. As the SAM mutant retains the predicted propeptide cleavage site RQKR, mutant Dsg1 that still gets into the endoplasmic reticulum (ER) is likely to be processed correctly in the Golgi. To test this idea, we first assessed the extent to which Dsg1 missing 12 amino acids encoded by Exon 2 (SAM-Dsg1), mimicking the c.49-1 G>A mutation, accumulated on the plasma membrane. The SAM mutant exhibited decreased plasma membrane localization with a corresponding increase in cytoplasmic staining in primary keratinocytes (Figure 6A, B), recapitulating what has been reported for this mutant in patient tissues (Cohen-Barak et al., 2020; Samuelov et al., 2013).

**Figure 6.**
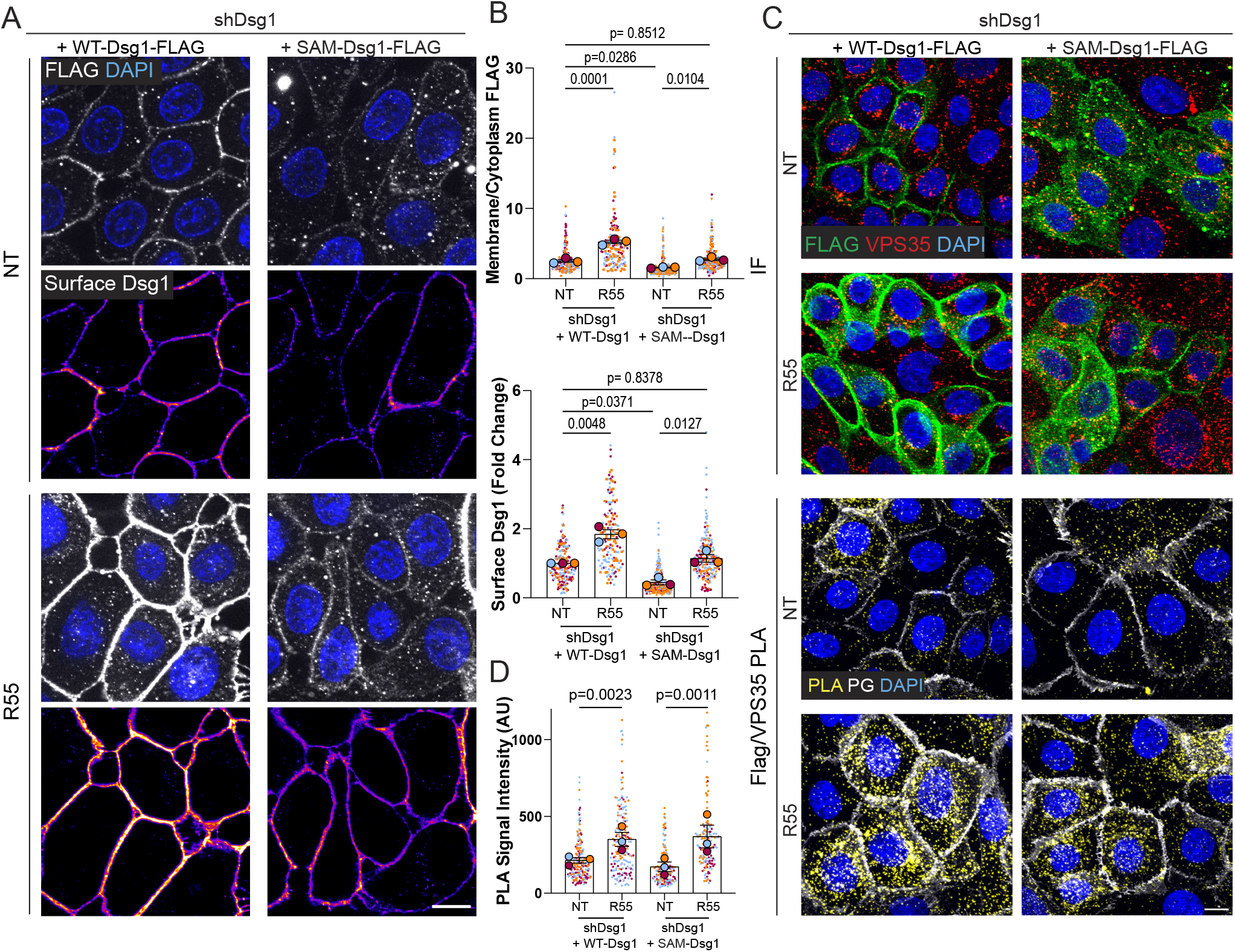
Retromer function-enhancing chaperone, R55, increases WT- and SAM-Dsg1 plasma membrane localization and proximity to VPS35. (A) Untreated (NT) and R55-treated cells expressing either a FLAG-tagged WT- or SAM-Dsg1 construct were surface-labeled for Dsg1 in live cells with anti-Dsg1 antibody. Cells were also fixed, permeabilized, and stained with an anti-FLAG antibody. Scale bar, 10 µm. (B) Quantification of average surface Dsg1 and FLAG membrane/cytoplasmic ratio following R55 treatment; one-way ANOVA with Sidak post-hoc from three biological repeats, and error bars are SEM. (C) Top: Immunofluorescence staining (IF) using FLAG and VPS35 antibodies was done to confirm antibody recognition of the proteins of interest. Bottom: Tandem PLA using FLAG antibody paired with VPS35 antibody to test VPS35 proximity to FLAG-tagged WT- and SAM-Dsg1 with and without R55 treatment. Plakoglobin (PG) staining is included to label cell-cell membranes (white). Scale bar, 10 µm. (D) Quantification of FLAG-VPS35 PLA +/- R55; two-way ANOVA with Sidak post-hoc test from three biological repeats, and error bars are SEM.

Because the retromer is necessary for Dsg1 exit from the endo-lysosomal system, we wanted to test if enhancing retromer function could increase SAM-Dsg1 plasma membrane localization. A small molecule chaperone called R55 was developed to enhance the stability and function of the retromer (Mecozzi et al., 2014). To test if enhancing retromer stability resulted in increased VPS35 association with Dsg1 in keratinocytes expressing FLAG-tagged WT-Dsg1 or the SAM-Dsg1 mutant, PLA was performed using a VPS35-FLAG antibody pair. R55 enhanced VPS35 association with both WT- and SAM-Dsg1 (Figure 6C, D). Although there was no noticeable change in Dsg1 protein levels with the addition of R55, the chaperone enhanced WT- and SAM-Dsg1 localization on the plasma membrane, such that the level of SAM-Dsg1 on the plasma membrane was now comparable with WT-Dsg1 (Figure S5C, D and Figure 6A, B). These data illustrate that the SAM mutant can undergo retromer-mediated trafficking, and that R55 is sufficient to enhance WT- and SAM-Dsg1 on the plasma membrane.

### The retromer chaperone, R55 enhances Dsg1-dependent keratinocyte stratification

In light of R55’s ability to increase cell surface Dsg1, we hypothesized that this retromer chaperone would enhance keratinocyte stratification (Figure 5F, G). To test this idea, GFP labeled keratinocytes were pre-treated with R55 for 24hrs before mixing in a 50:50 ratio with unlabeled keratinocytes and R55 treatment was continued for the next 24 hours during which time stratification occurred. As expected, shRNA-mediated depletion of Dsg1 resulted in decreased stratification (Figure 7A, B). Addition of R55 enhanced the stratification of control GFP-labeled keratinocytes, but not Dsg1-depleted keratinocytes (Figure 7A, B). Ectopic expression of WT-Dsg1, which forces Dsg1 expression earlier than in unlabeled cells expressing endogenous Dsg1, rescued stratification of Dsg1-depleted keratinocytes above normal levels. R55 did not further enhance the number of stratifying cells under this condition, likely as an upper limit or threshold was reached in cells ectopically expressing WT-Dsg1 (Figure 7A, B). Importantly, while the SAM mutant rescued stratification to a certain extent in untreated cells, R55 significantly increased stratification up to or beyond levels induced by WT-Dsg1 (Figure 7A, B). Overall, these data illustrate that R55 can enhance Dsg1-dependent stratification of keratinocytes including cells expressing the trafficking-deficient SAM-associated mutation (Figure 7C).

**Figure 7.**
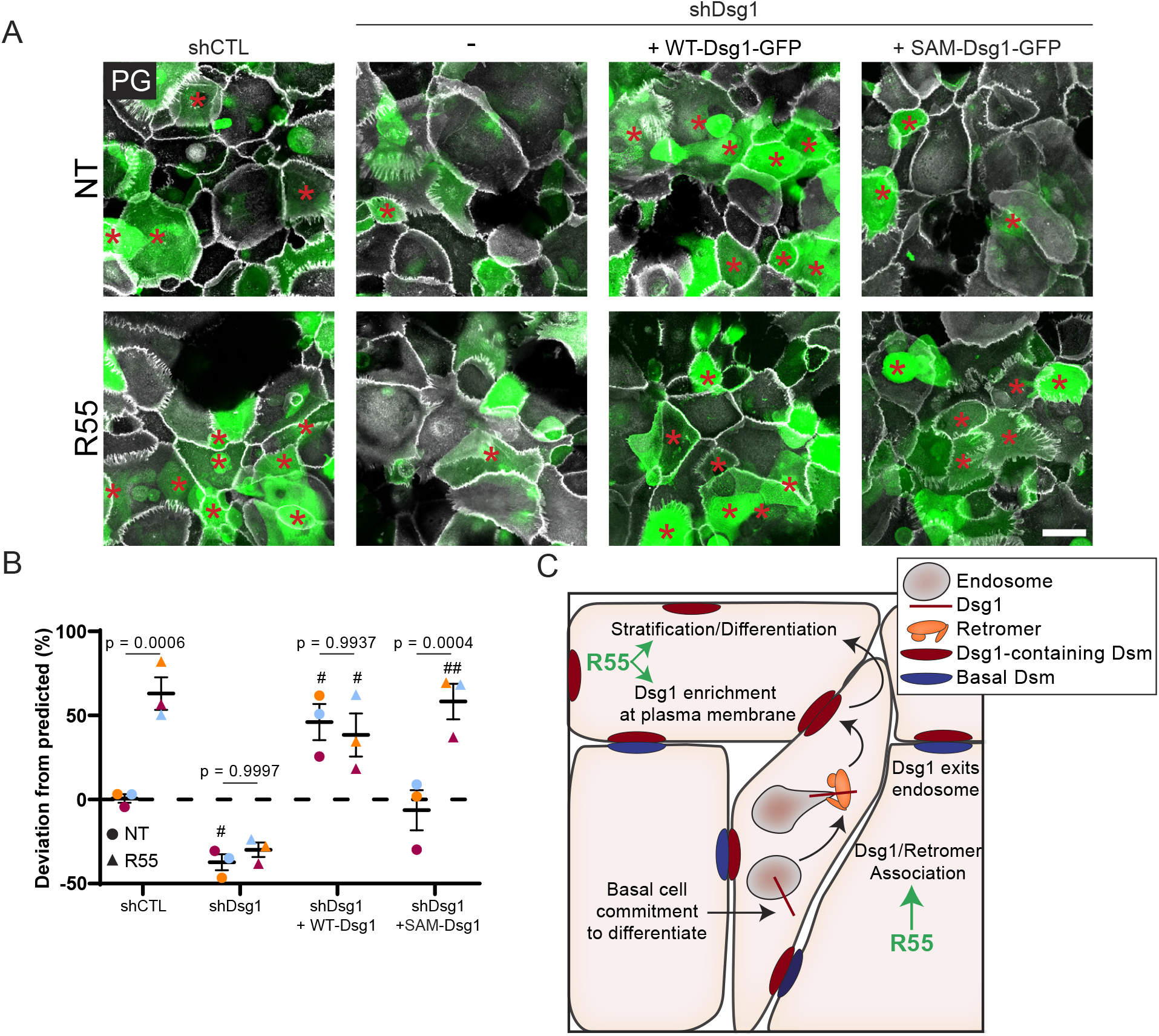
R55 enhances keratinocyte stratification in cells expressing endogenous Dsg1 and SAM-Dsg1. (A) Keratinocytes expressing shCTL, shDsg1, shDsg1+ WT-Dsg1-GFP, or shDsg1+SAM-Dsg1-GFP were pretreated +/- R55 for 24hrs, mixed 50:50 with unlabeled WT cells and allowed to settle overnight in 0.07mM CaCl_2_ media +/- R55. Cells were switched to 1.2mM CaCl_2_ media +/- R55 for 24hrs then fixed and stained with Plakoglobin (PG)/DAPI. Scale bar, 50 µm. (B) Quantification of deviation from predicted % of GFP cells in the stratified layer; one-way ANOVA with Tukey post-hoc test from three biological repeats, and error bars are SEM. When comparing NT and R55 treatments for each condition, p-values are shown. When comparing conditions to the control (shCTL), significant p-value <0.05 or <0.01 denoted with # or ##, respectively. (C) Model illustrating our findings that the retromer promotes Dsg1 plasma membrane localization to promote epidermal differentiation processes such as stratification. The addition of the retromer enhancing chaperone, R55, enhances WT- and SAM-Dsg1 association with the retromer, enriches WT- and SAM-Dsg1 on the plasma membrane, and enhances stratification of cells expressing endogenous Dsg1 and SAM-Dsg1.

## DISCUSSION

Intracellular trafficking coordinates with changes in the cytoskeleton and cell polarity machinery to regulate developmental and morphogenetic processes as well as tissue homeostasis (Xie et al., 2018). Alterations in protein trafficking and consequent defects in cell polarity are known to impair barrier and transport functions in simple epithelia. Interfering with basolateral sorting, endocytosis, recycling and transcytosis can lead to diseases such as cystic fibrosis, Wilson disease, familial hypercholesterolemia and ADPKD (Charron et al., 2000; Forbes and Cox, 2000; Koivisto et al., 2001; Moyer et al., 1999; Stein et al., 2002). Less is known about the importance of intracellular trafficking in polarized multi-layered epithelial such as the epidermis. Here, we identify a role for the retromer trafficking machinery in regeneration of the multi-layered epidermis. We show that the retromer functions at least in part by its selective association with and recycling of desmosomal cadherin Dsg1, to drive epidermal stratification and differentiation. We also show that the retromer can be exploited to enhance delivery, and consequently, function of a Dsg1 mutant associated with a severe, systemic disorder called SAM syndrome (Figure 7C).

The retromer was originally discovered in yeast to be required for retrieval of a sorting receptor called Vps10 from the endosome to the TGN (Seaman et al. 1998), and is now well-established to be more broadly involved in processes of endosomal trafficking and recycling. Since its discovery, the retromer has been shown to play essential roles in specific aspects of development, such as wing morphogenesis in flies, and establishing anterior/posterior neuronal polarity in *C. elegans* (Belenkaya et al., 2008; Franch-Marro et al., 2008; Port et al., 2008; Prasad and Clark, 2006; Yang et al., 2008). While a study in HeLa cells identified ∼150 retromer cargoes (Steinberg et al., 2013), in vivo, the retromer seems to be involved in selected pathways, for instance the Wnt, but not Hh, Notch or BMP pathways in Drosophila wing development (Belenkaya et al., 2008; Franch-Marro et al., 2008; Port et al., 2008). In mammals, depletion of retromer components affects specific aspects of early development, leading to embryonic lethality between E8.5-10 embryos (Radice et al., 1991; Wen et al., 2011). In humans, specific roles for the mammalian retromer are emerging due to the association of the complex with diseases such as ALS, Alzheimer’s and Parkinson’s disease. In complex epithelia, the retromer has been associated with HPV infection. However, its role in normal homeostasis of complex epithelia such as the epidermis has not been addressed.

Our data show that the endosomal trafficking retromer complex promotes endosomal trafficking of the stratified tissue-specific desmosomal cadherin Dsg1, but not the AJ cadherin E-cadherin, in epidermal keratinocytes. Although retromer-mediated trafficking seems to be specific to Dsg1 when compared to E-cadherin in keratinocytes, it has been reported that the retromer component Snx1 affected bulk-internalized E-cadherin recycling in simple epithelial cells (Bryant et al., 2007). However, the authors were unable to find an interaction between E-cadherin and Snx1 and suggested that the effect on E-cadherin internalization is indirect, consistent with our finding that E-cadherin doesn’t require retromer-mediated trafficking in steady state conditions. The specificity of the retromer machinery for Dsg1 and not E-cadherin could help explain how these proteins are segregated into different domains during keratinocyte differentiation, to promote actin reorganization and tension redistribution required for delamination (Nekrasova et al., 2018). Delamination is an important contributor to the process of stratification in conjunction with the asymmetric division of basal cells to establish and maintain the epidermal layer (Damen et al., 2021; Nekrasova et al., 2018; Williams et al., 2014). The dynein light chain Tctex-1 was also found to be necessary for Dsg1’s proper localization on the insoluble plasma membrane, segregated from E-cadherin (Nekrasova et al., 2018). As dynein has been shown to interact with the retromer to promote endosome-to-TGN trafficking (Hong et al., 2009; Wassmer et al., 2009), it seems plausible that the retromer works with dynein/Tctex-1 to tailor Dsg1-specific trafficking to promote proper plasma membrane localization required for Dsg1’s functions in morphogenesis.

While not addressed in this study, it is also possible that the retromer contributes to later stages of differentiation by regulating the exchange of suprabasal desmosomal cadherins into desmosomes containing basal cadherins, during a cell’s transit into superficial layers. For instance, suprabasal Desmocollin 1 becomes increasingly incorporated into desmosomes expressing Desmocollin 3 as cells transit into superficial layers (North et al., 1996). The mechanism of this exchange is not clear but is likely important for establishing the patterned distribution of desmosomal cadherins, which is functionally relevant. For example, during epidermal morphogenesis, the stratified tissue-specific Dsg1 regulates multiple processes important for maintaining epidermal homeostasis, including patterning the function of EGFR and ErbB2 to promote the basal to suprabasal transition while maintaining the tight junction barrier in the superficial layer (Broussard et al., 2021; Getsios et al., 2009).

Lastly, this work identified the retromer as a target for therapeutic intervention in skin disease. Previous work established that the retromer is necessary for HPV infection and identified a HPV specific peptide that inhibits the association of the retromer with the cellular cargo, DMT1-II (Carosi et al., 2021; Lipovsky et al., 2013; Zhang et al., 2020). However, most investigation into the retromer’s role in human health has been focused on neurological diseases and tauopathies such as Parkinson’s disease, Alzheimer’s disease, and ALS (Carosi et al., 2021). Enhancement of retromer activity by the retromer chaperones R55 and R33 has been shown to decrease pathogenic amyloid-precursor protein (APP) processing and tau phosphorylation in cultured mouse hippocampal neurons or human iPSC-derived neurons (Mecozzi et al., 2014; Young et al., 2018). Furthermore, treatment of animal models for Alzheimer’s disease and ALS with R33 or an R55-derivative, Compound 2a, ameliorated memory impairment, APP processing, tau phosphorylation and neuroinflammation or attenuated lysosomal defects and Golgi fragmentation, respectively (Li et al., 2020; Muzio et al., 2020). Here we showed that treatment of keratinocytes with R55 enhanced epidermal stratification by bolstering the trafficking of either WT-Dsg1 or a SAM syndrome-associated Dsg1 mutant, illustrating a new role for the retromer as a therapeutic target for epidermal diseases. Additionally, as Dsg1 is also disrupted in common disorders such as psoriasis, squamous cell carcinoma, and eosinophilic esophagitis, retromer therapy may have broad benefit beyond rare genetic disorders (Godsel et al., 2020; Hegazy et al., 2021; Sherrill et al., 2014; Wong et al., 2008).

Collectively, our observations provide an alternative perspective to the textbook view of desmosomes as stable, adhesive spot welds, supporting the emerging concept that these intercellular junctions function as dynamic complexes that undergo constant remodeling during epidermal regeneration and adaptation to environmental and mechanical stimuli (Hegazy et al., 2021; Kottke et al., 2006; Müller et al., 2021). They also reveal the importance of understanding Dsg1 trafficking mechanisms for determining underlying molecular mechanisms, and potentially treating, disorders of stratified epithelia.

## Acknowledgments

Research was supported by NIH/NIAMS P30 AR075049 awarded to Northwestern University Skin Biology & Diseases Resource-Based Center. Imaging work was performed at the Northwestern University Center for Advanced Microscopy generously supported by NCI CCSG P30 CA060553 awarded to the Robert H Lurie Comprehensive Cancer Center. Histology services were provided by the Northwestern University Research Histology and Phenotyping Laboratory which is supported by NCI P30-CA060553 awarded to the Robert H Lurie Comprehensive Cancer Center. This work was supported by NIH/NIAMS R01 AR041836, NIH/NIAMS R01 AR043380, and NIH/NCI R01 CA228196 with partial support from J.L. Mayberry Endowment to KJG. MH was supported by NIH/NCI T32 CA009560 and NIH/NIAMS F31 AR076188. JAB was supported by NIH/NIAMS K01 AR075087 and NIH/NIAMS T32 AR060710.

## MATERIAL AND METHODS

### Cell culture and drug treatments

Primary keratinocytes were isolated as previously described from human neonatal foreskin obtained from Northwestern University’s Skin Biology and Disease Resource-based Center (Arnette et al., 2016). Undifferentiated keratinocytes were cultured in M154 media (Gibco) supplemented with 0.07mM CaCl_2_, human keratinocyte growth supplement (HKGS), gentamicin, and amphotericin B. To induce differentiation in 2D cultures, the calcium concentration of the growth medium was raised to 1.2mM CaCl_2_. 3D organotypic epidermal raft cultures using primary keratinocytes were generated as previous described (Arnette et al., 2016). Drug treatments of 500μM Primaquine for 6hr-overnight and 10μM R55 for 48-72hrs were the conditions used. Primaquine treatment of pre-differentiated keratinocytes followed 1.5 days in 1.2mM CaCl_2_ medium. During time-lapse imaging, keratinocytes were cultured with FluoroBrite DMEM (Gibco).

### Antibodies and fluorescent dyes

The following primary antibodies were used for this study: rabbit anti-VPS35 (Abcam cat# ab157220, RRID:AB_2636885; Life Technologies cat# PA5-21898, RRID:AB_11153540), mouse anti-VPS35 (Santa Cruz Biotechnology cat# sc-374372, RRID:AB_10988942), mouse anti-VPS29 (Santa Cruz Biotechnology cat# sc-398874, RRID:AB_2885036), mouse anti-Dsg1 (P124 - PROGEN cat# 651111, RRID:AB_1541107; R&D Systems cat# MAB944, RRID:AB_2093296; 27B2 – Thermo Fisher Scientific cat# 32-6000, RRID:AB_2533088), rabbit anti-Dsg1 (Abcam cat# ab124798, RRID:AB_10974963), mouse anti-FLAG (M2; Sigma-Aldrich Cat# F1804, RRID:AB_262044), chicken anti-plakoglobin (1407 and 1408 - Aves Laboratories), mouse anti-EEA1 (BD Bioscience cat# 610456, RRID:AB_397829), mouse anti-LAMP1 (Santa Cruz Biotechnology cat# sc-20011, RRID:AB_626853), mouse anti-GFP (JL8 - Takara cat# 632381, RRID: AB_2313808), mouse anti-E-cadherin (HecD1-provided by M. Takeichi and O. Abe, Riken Center for Developmental Biology, Kobe, Japan), rabbit anti-E-cadherin (795-provided by R. Marsh and R. Brackenbury, University of Cincinnati, Cincinnati, OH), goat anti-E-cadherin (R&D Systems cat# AF748, RRID:AB_355568), rabbit anti-β-catenin (Sigma cat# C2206, RRID:AB_476831), rabbit anti-K10 (provided by J. Segre, National Human Genome Research Institute, Bethesda, Maryland, USA), rabbit anti-M6PR (cation independent) (Abcam cat# ab124767, RRID:AB_10974087), rabbit anti-GLUT1 (Abcam cat# ab115730, RRID:AB_10903230), rabbit anti-GAPDH (Sigma-Aldrich cat# G9545, RRID: AB_796208), mouse anti-tubulin (12G10; Developmental Hybridoma Studies Bank, RRID: AB_1157911), mouse anti-actin (C4; Millipore Sigma, cat# MAB1501, RRID:AB_2223041), mIgG isotype control (Abcam, cat# ab37355, RRID:AB_2665484), rbIgG isotype control (Abcam, cat# ab27478, RRID:AB_2616600).

Secondary antibodies and dyes used for immunofluorescent studied include goat anti-mouse, - rabbit, and -chicken conjugated to Alexa Fluor (AF) fluorophores (Thermo Fisher Scientific), AF-conjugated goat anti-mouse IgG1 (Thermo Fisher Scientific), AF-conjugated goat anti-mouse IgG2a (Thermo Fisher Scientific), AF-conjugated donkey anti-mouse (Thermo Fisher Scientific), unconjugated goat anti-mouse (Jackson Laboratory). To stain nuclei, 4′,6-Diamido-2-Phenylindole (DAPI, Sigma-Aldrich cat # D9542) was used. LysoBrite stained for acidic endo-lysosomal organelles (AAT Bioquest cat# 22643). Secondary antibodies used for immunoblotting include HRP conjugated goat anti-mouse and -rabbit (Kirkegaard Perry Labs) and NIR-fluorescent IRDye 680 or 800 goat anti-mouse IgG, -mouse IgG1, -mouse IgG2a, - rabbit (Licor).

### siRNA-mediated transfection

Keratinocytes were transiently transfected with siGENOME non-targeting siRNA pool #2 (Dharmacon cat# D-001206-14), siRNA against VPS35 from Sigma Silencer Select with a targeting sequence 5’-CCATGGATTTTGTACTGCTTT-3’ and Integrated DNA Technologies with a targeting sequence 5’-GAGATATTCAATAAGCTCAACCTTG-3’, siRNA against VPS29 (Integrated DNA Technologies) with a targeting sequence 5’-ATGATGTGAAAGTAGAACGAATCGA3’, siRNA directed against Dsg1 (Integrated DNA Technologies) with a targeting sequence 5′-CCATTAGAGAGTGGCAATAGGATGA-3′ using Amaxa Nucleofector System (Lonza) for electroporation of cells according to manufacturer’s instructions.

### Plasmids and viral transduction

Generation of pLZRS-GFP, pLZRS-WT-Dsg1-FLAG, and pLZRS-shDsg were generated by Getsios et al. (Getsios et al., 2009). pLZRS-SAM Dsg1-FLAG (deletion of Exon 2) was generated as described in Cohen-Barak et al. (Cohen-Barak et al., 2020). pLZRS-shNT (Non-targeting) was generated as described (Arnette et al., 2020). Silencing resistant pLZRS-WT-Dsg1-FLAG/GFP and pLZRS-SAM Dsg1-FLAG/GFP were newly generated (Epoch Life Science Inc.). pLZRS constructs were transiently transfected into Phoenix cells using Lipofectamine 2000. Phoenix cells that were successfully transfected with the plasmid were selected using 1 μg/ml puromycin. Following a 24-48hr incubation at 32°C, retrovirus-containing Phoenix cell supernatant was collected and concentrated using Amicon Ultra-15 Centrifugal Filter Unit (EMD Millipore). Keratinocytes were infected with concentrated retrovirus diluted in growth medium and 4μg/ml polybrene at 20% confluency for 1.5 hours at 32°C.

### Immunoblotting and immunoprecipitation

Keratinocytes were lysed using Urea Sample Buffer (8 M deionized urea, 1% SDS, 60 mM Tris, pH 6.8, 10% glycerol, 0.1% pyronin-Y, 5% β-mercaptoethanol). Proteins were separated using SDS-PAGE electrophoresis before transfer to a nitrocellulose membrane (0.2-0.45 μm pore size). Membranes were blocked with either 5% Milk or LICOR Intercept (PBS) Blocking Buffer, (Licor). Primary and secondary antibodies were diluted in blocking buffer. Immunoreactive proteins were visualized using NIR fluorescence or chemiluminescence using the Odyssey Imaging system (Licor). When probing for VPS29, lysates were run on a gradient gel. Following electrophoresis, the gel was cut in with the upper half subjected to a normal transfer protocol and the lower half undergoing a quick transfer (80V for 40min) on nitrocellulose membrane (0.2 μm pore size).

For immunoprecipitation, keratinocytes infected with WT-Dsg1 FLAG virus were grown to confluency before being differentiated overnight with 1.2mM Ca^2+^ media. Cells were then lysed and scraped in ice cold buffer (50mM Tris-HCl pH 7.5, 1mM EDTA pH 8, 0.5% Triton X-100, 150mM NaCl, and 10% glycerol) and transferred to 1.5 ml tubes. Following 1min vortex the lysate was allowed to sit on ice for 10 min followed by centrifugation at 4°C for 30 min at 21,130 x g. 5% volume of the supernatant was saved as the input, the remaining supernatant incubated with either rabbit-IgG primary antibody (negative control) and VPS35 antibody (Life Technologies). Following primary antibody incubation, Dynabeads Protein G (Life Technologies) were used per manufacturer’s protocol. IP samples were eluted using 3X Lammli sample buffer and boiled before proceeding to standard immunoblotting protocol.

### Immunofluorescence and histology

Primary keratinocytes were cultured on 12-15mm glass coverslips. Cells were fixed with either 4% paraformaldehyde solution for 10-15min room temperature or 100% ice-cold anhydrous methanol for 2min on ice. For live cell staining of surface proteins, primary antibody was added before fixation for 1-2hr(s) on ice. To stain for cytoplasmic domains of transmembrane proteins or cytoplasmic proteins in PFA-fixed cells, 0.2% Triton X-100 was used to permeabilize the cells for 10 min at room temperature. Cells were then blocked with 1% BSA solution for 30 minutes at room temperature. 5% normal goat serum was added to the blocking solution when primary antibodies did not include goat IgG. Primary antibodies and secondary antibodies were diluted in block solution and added to cells for either 1-2hrs at 37°C or overnight at 4°C. Paraffin-embedded tissue sections of 3D organotypic rafts were baked at 60°C overnight. Xylene/ethanol was used to deparaffinize the tissue section and 0.5% Triton X-100 permeabilized the cells. 0.01M citrate buffer was used for antigen retrieval. 2D and 3D samples were mounted using ProLong Gold antifade reagent to glass slides. 3D raft tissue sections were also processed for hematoxylin and eosin (H&E) staining using standard methods.

### Proximity ligation assay

Primary keratinocytes were plated on glass coverslips and fixed with 4% PFA for 10-15min and permeabilized with 0.2% Triton for 10min. Proximity ligation assay protocol, image acquisition and analysis were performed as previously described (Hegazy et al., 2020).

### Image acquisition and analysis

Apotome images were acquired with an epifluorescence microscope (Axio Imager Z2, Carl Zeiss) equipped with an Apotome.2 slide module, Axiocam 503 Mono digital camera, X-Cite 120 LED Boost System, and Plan-Apochromat 40x/1.4 and 63x/1.4 objective. H&E images were obtained with a microscope (DMR, Leica) equipped with 10x and 40x objective a digital camera (Leica DFC295) using Leica Application Suite software. Confocal images were acquired with a Nikon A1R+ confocal with GaAsP detectors and Nikon W1 Spinning Disk Confocal using NIS Elements software (Nikon) and equipped with 95B prime Photometrics camera, Plan-Apochromat 60x/1.4 objective and Plan-Apochromat 40x/1.3 objective. Time lapse imaging with Nikon A1R+ was performed using Plan-Apochromat TIRF 100x/1.49 objective and captured with no delay. To improve image quality, background in A1R confocal images was subtracted using rolling ball background subtraction followed by 2D deconvolution using NIS Element software (Nikon). 3D reconstruction of A1R confocal z-stack images was implemented with Volume Viewer’s Depth Coded Alpha Blending (rainbow contrast look up table).

### Antibody-based recycling assay

Keratinocytes transfected with siCTL, siVPS35, or siVPS29 were each plated on three glass coverslips. All three coverslips were incubated with anti-Dsg1 monoclonal antibody for 1hr on ice. Following PBS washes, two coverslips were switch to 0.07mM low calcium media with 3mM calcium chelator EGTA for 30min at 37°C to promote endocytosis while the third coverslip remained on ice (Steady State). Following internalization and subsequent PBS washes, unconjugated goat anti-mouse secondary was added to the two coverslips on ice to block remaining surface labeled Dsg1. 1.2mM high calcium media was added to one of the PBS-washed coverslips and incubated in 37°C for 30min to induce recycling while the second coverslip remained on ice (Block/Internalization). Following 30min incubation, the last coverslip was PBS washed on ice (Recycle). To stain for surface Dsg1 (Steady State), residual Dsg1 (Block/Internalization), and recycled Dsg1 (Recycle), AF488-conjugated Goat anti-mouse secondary antibody was added the live cells for 30min before cells were fixed with 4% PFA and permeabilized with 0.1-0.2% Triton-X 100 for 10min. To stain for the internalized pool AF568-conguated goat anti-mouse was added following permeabilization. Internalization of Dsg1 was quantified using the 568 channels in the Block/Internalization coverslip and normalized to intensity of Dsg1 at Steady State. Residual Dsg1 was quantified using the intensity of 488 channel of the Block/Internalization coverslip and normalized to the internalization pool in the same coverslip. The recycling pool of Dsg1 was quantified using the intensity of 488 channel of the Recycle coverslip and normalized for both internalized signal in the same coverslip and residual Dsg1.

### Stratification assay

Stratification assays and analysis were performed as previously described for Dsg1-GFP stratification studies (Broussard et al., 2021), allowing GFP/Dsg1-GFP labeled keratinocytes mixed with unlabeled keratinocyte to stratify for 24hrs. For retromer depletion, VPS35 and VPS29 siRNA were transfected into undifferentiated GFP/Dsg1-GFP expressing keratinocytes before mixing with unlabeled keratinocytes in a known ratio. For R55 stratification experiments, GFP-labeled keratinocytes expressing shCTL, shDsg1, shDsg1+WT-Dsg1, and shDsg1+SAM-Dsg1, were pretreated for 24hrs before mixing with unlabeled keratinocytes with continued R55 treatment. Following 24hrs of stratification, 4% paraformaldehyde-fixed cells were processed as described above and stained for plakoglobin and DAPI. Confocal images with 0.5μm z-stacks were obtained to calculate the ratio of stratified cells that are GFP or Dsg1-GFP positive. The percentage of GFP-positive stratified cells per condition was compared to the predicted percentage of cells that express GFP after mixing with unlabeled cells to obtain the percent deviation from predicted,

### Quantification and statistical analysis

A minimum of three independent experiments (biological repeats) was performed for each experiment. For immunofluorescence studies, population analysis of each independent experiment was performed with a minimum of 15-30 cells spanning at least 3 areas on the coverslip. All images in each experiment were captured using the same parameters. Fluorescence intensity, area, aspect ratio, and roundness was analyzed using FIJI/ImageJ software (NIH). For biochemical assays, bands on the immunoblots were quantified using LiCOR Image Studio (Version 5.2) software. Graphing and statistical analyses were done on GraphPad Prism (version 8.0) software. Specific tests are indicated in the figure legend. The graphs show the mean of the independent experiments represented with large circles as well as the small individual data points with colors corresponding to each independent experiment when applicable. The error bars represent standard error of the mean (SEM).

## Notes

### Competing Interest Statement

The authors have declared no competing interest.

