## Supplemental Figures for "Epidermal Stratification Requires Retromer-Mediated Desmoglein-1 Recycling"

**Movie 1. Retromer depletion disrupts Dsg1-GFP localization in intracellular tubulovesicular structures**. Representative time-lapse images of Dsg1-GFP in keratinocytes that were differentiated overnight in cells transfected with siCTL, siVPS35, or siVPS29. Yellow arrowheads illustrate Dsg1 localization in vesicle tubules and projections. Images were pseudocolored based on signal intensity using the Fire lookup table. Interval time between frames is 2.15 seconds. Movie corresponds with images in Figure 3C. Scale bars, 10 µm, 1µm.

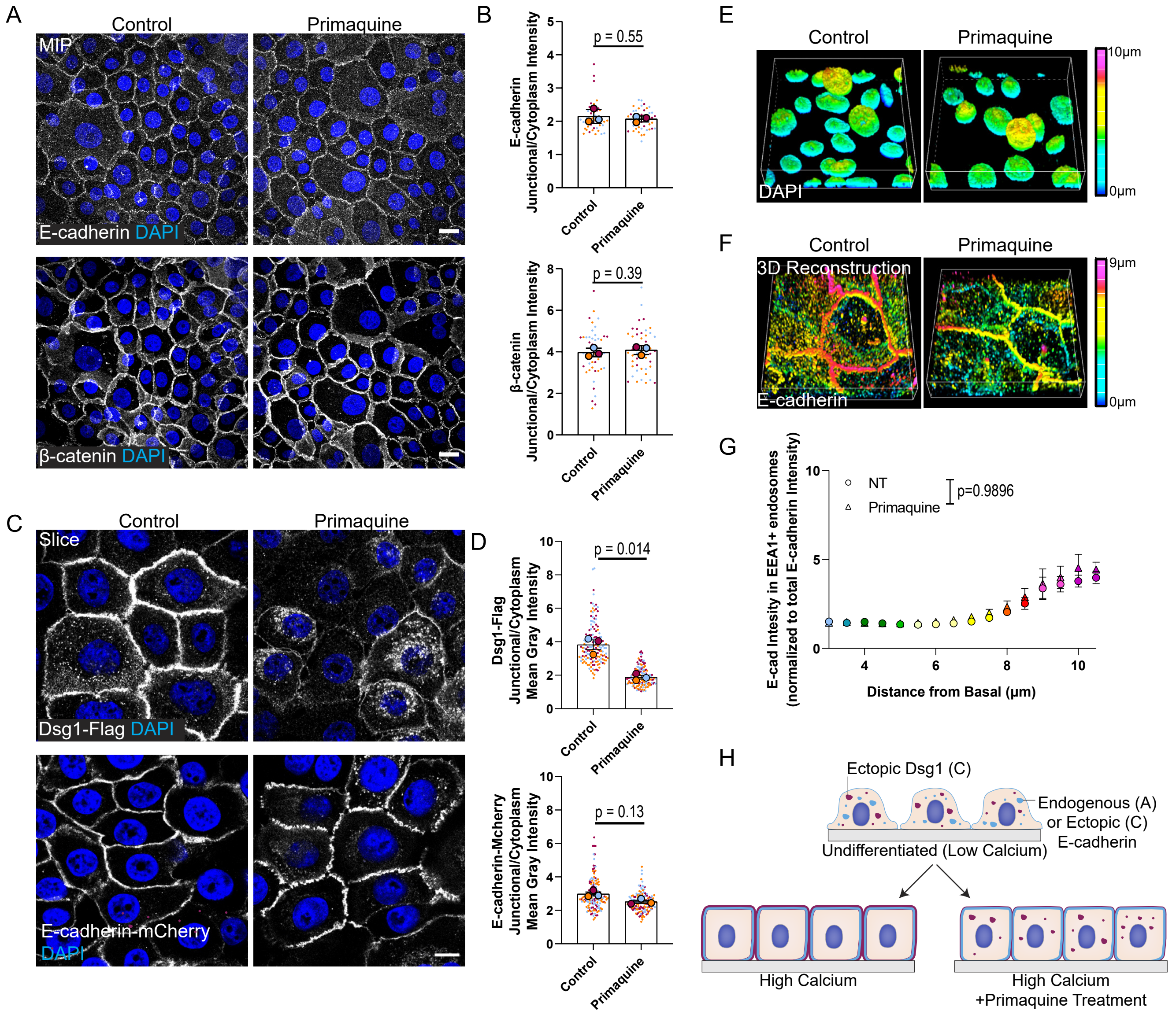


**Figure S1. Primaquine treatment disrupts ectopic Dsg1, but not AJ proteins.** (A) Maximum intensity projection (MIP) of E-cadherin and β-catenin immunofluorescence following overnight Primaquine treatment (200µM) of undifferentiated keratinocytes. Scale bar, 20 µm. (B) Quantification of E-cadherin and β-catenin membrane/cytoplasmic ratio following Primaquine treatment; paired t-test from three biological repeats, and error bars are SEM. (C) Dsg1-FLAG and E-cadherin-mCherry immunofluorescence following overnight Primaquine treatment (200µM) of undifferentiated keratinocytes. Scale bar, 10 µm. (D) Quantification of Dsg1-FLAG and E-cadherin-mCherry membrane/cytoplasmic ratio following Primaquine treatment; paired t-test from three biological repeats, and error bars are SEM. (E) 3D reconstruction of DAPI image co-stained with Dsg1 (corresponding to Figure 1A) with a z-depth pseudocoloring illustration. (F) 3D reconstruction of E-cadherin staining (corresponding to Figure 1B) with a z-depth pseudocoloring illustration. (G) Quantification of E-cadherin intensity in EEA1 positive vesicles with respect to the distance from the basal layer. Colors correspond with the z-depth scale shown in F; two-way ANOVA with Sidak post-hoc test from three biological repeats, and error bars are SEM. (H) Schematic demonstrating the effect of Primaquine treatment on Endogenous E-cadherin (see panel A) and ectopic E-cadherin or Dsg1 (see panel C).

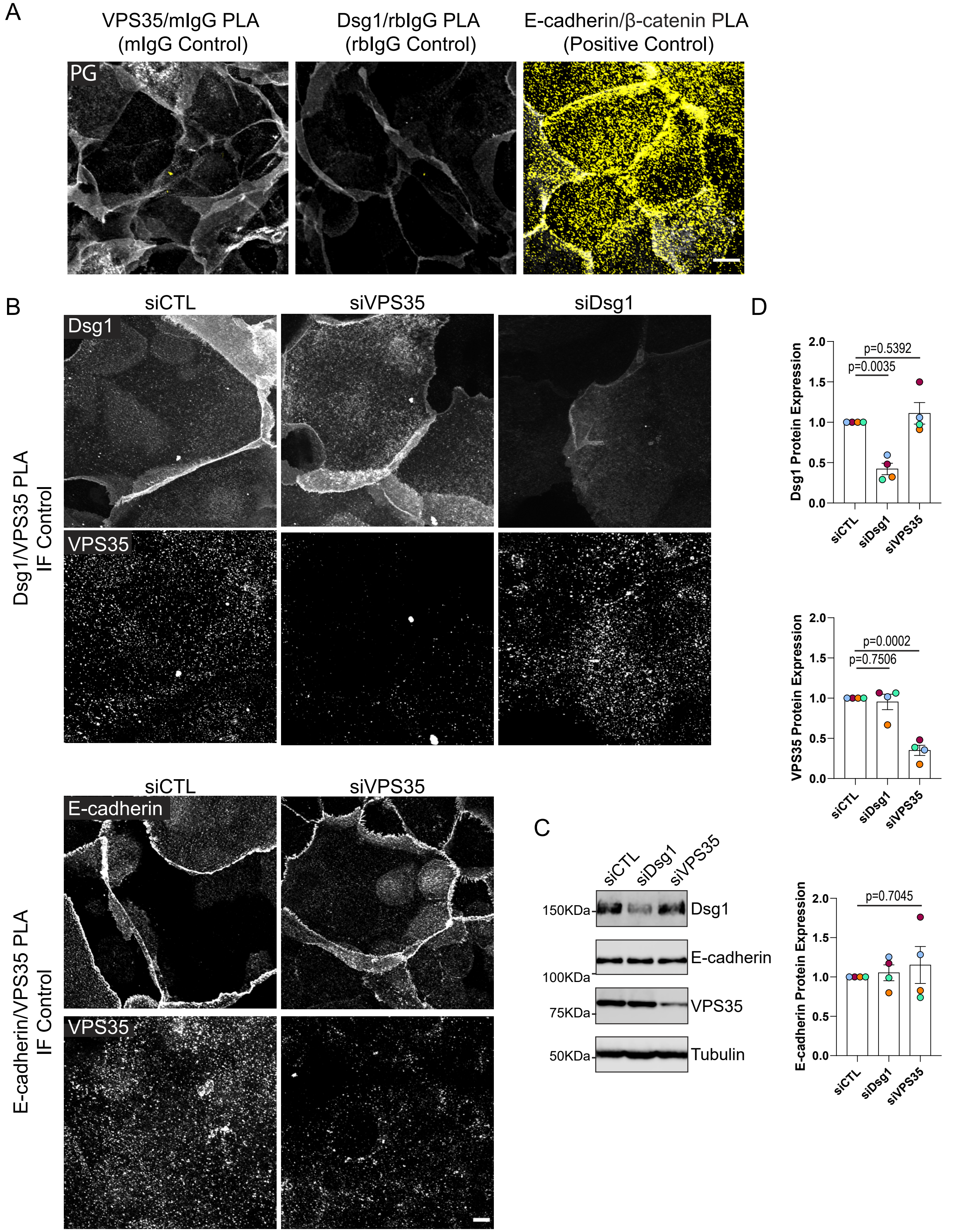


**Figure S2. Controls for the PLA used to assess the proximity of VPS35 to Dsg1 or E-cadherin.** (A) Non-specific interactions were tested by pairing anti-Dsg1 or anti-VPS35 antibodies with the species specific IgG corresponding to its partner's species: mouse IgG (mIgG Control), rabbit IgG (rIgG Control) and an E-cadherin/β-catenin antibody pairing used as a positive control for the PLA signal. Scale bar, 20 µm. (B). Immunofluorescence (IF) was done in tandem with PLA using the same antibody pairing at the same concentrations to confirm positive protein recognition by the antibodies. Scale bar, 10 µm. (C) Immunoblotting of lysates collected from cells grown in tandem with the PLA/IF coverslips to confirm efficient siRNA-mediated depletion. (D) Quantification of the densitometry of the immunoblot normalized to control levels; one-way ANOVA with Dunnett post-hoc test from three biological repeats, and error bars are SEM.

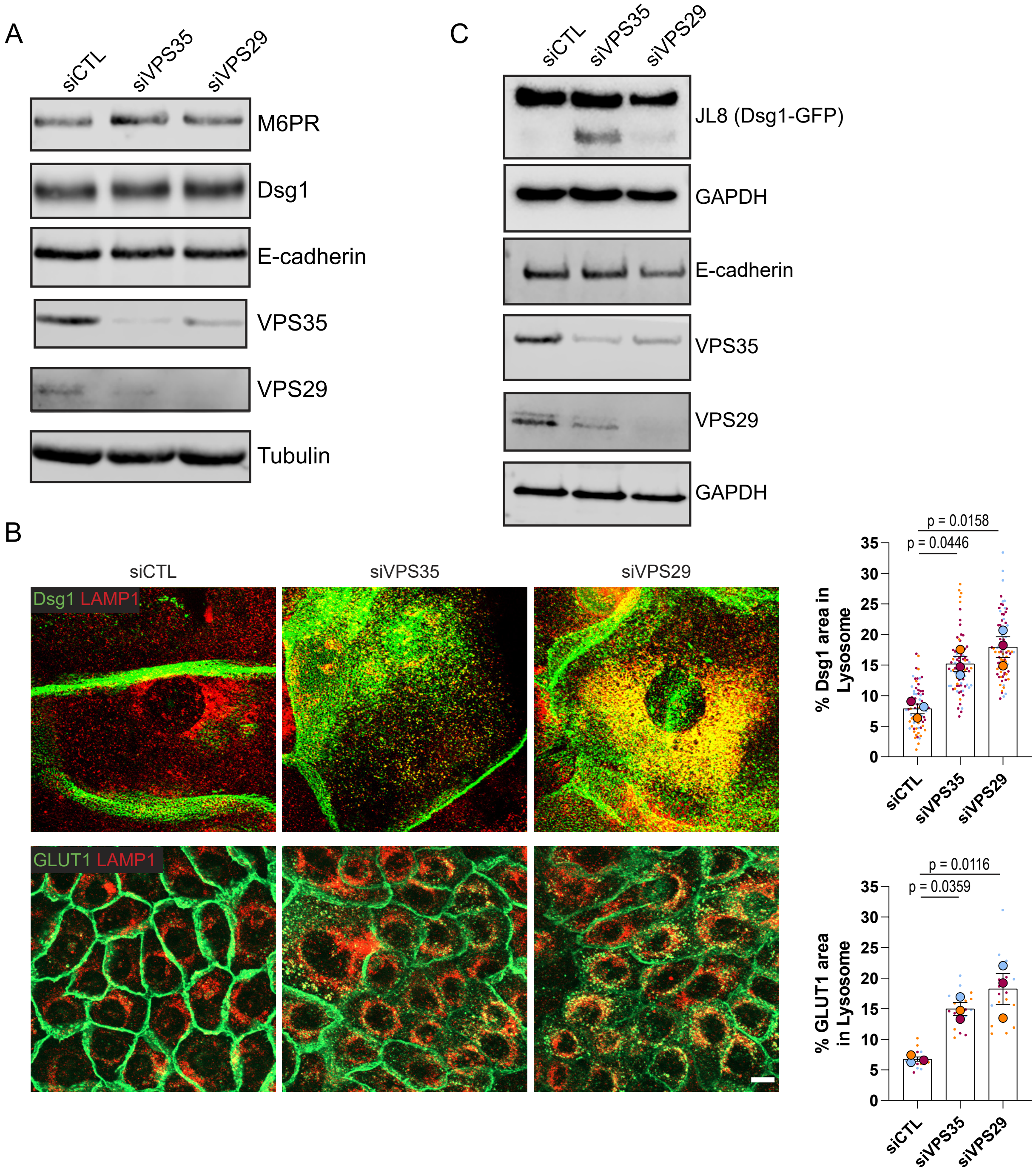


**Figure S3. Verification of retromer component depletion and endogenous Dsg1 colocalization with the endo-lysosomal marker LAMP1.** (A) Immunoblotting of lysates collected from cells grown in tandem with surface labeling of endogenous Dsg1 (see Figure 3A) and the antibody based recycling assay (see Figure 4) confirming efficient siRNA-mediated depletion of retromer protein targets. (B, Left) Immunofluorescence of endogenous Dsg1 in stratified cells and GLUT1, a known retromer cargo, in basal cells co-stained with the endo-lysosomal marker, LAMP1, in fixed cells following retromer component depletion. Scale bar, 10 µm. (B, Right) Quantification of the % Dsg1 area and % GLUT1 area in LAMP1 stained endo-lysosomal vesicles; one-way ANOVA with Dunnett post-hoc test from three biological repeats, and error bars are SEM. (C) Immunoblotting of lysates collected from cells grown in tandem with the Dsg1-GFP live cell imaging experiment (see Figure 3C, D).

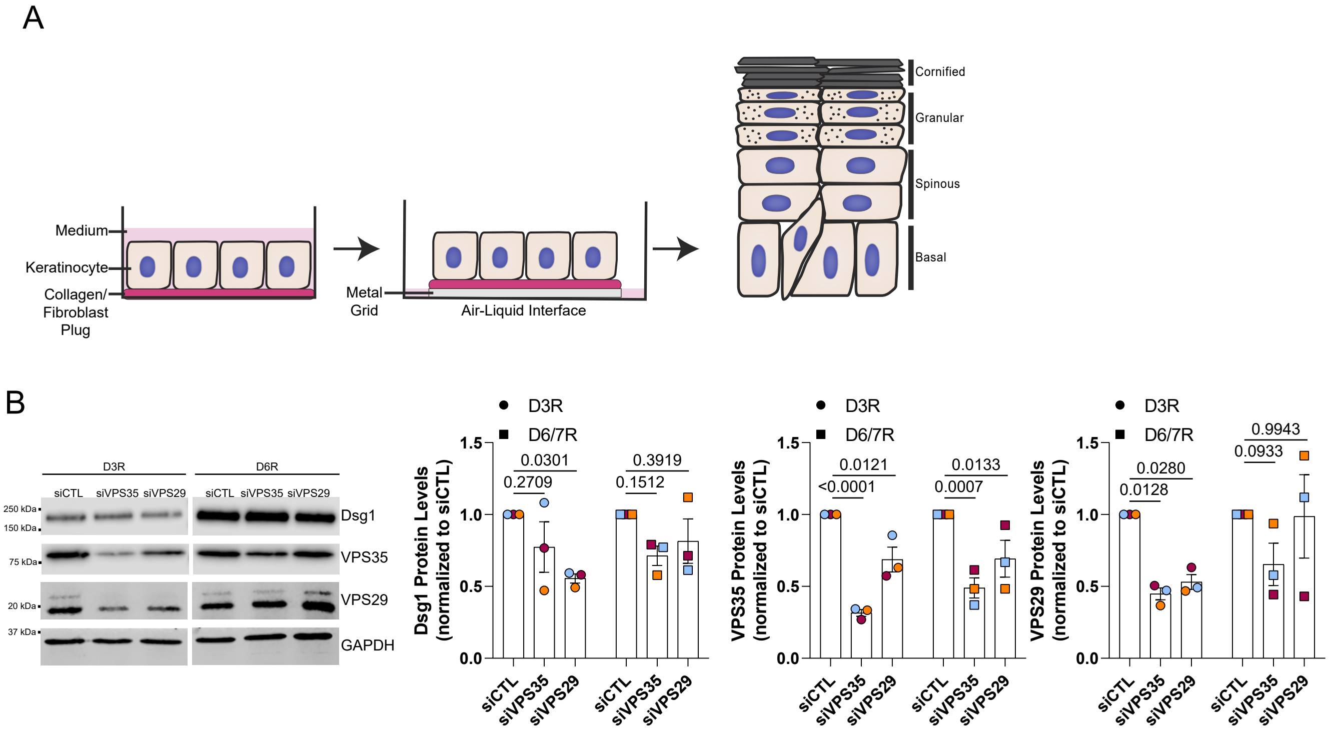


**Figure S4. 3D organotypic epidermal culture experimental design and immunoblots of day 3 and day 6-7 3D organotypic cultures.** (A) Keratinocytes were seeded on collagen plugs containing fibroblasts, which were subsequently lifted to an air-liquid interface by placing them onto a metal grid and feeding the culture from the bottom to induce stratification. Cultures were harvested 3 days or 6-7 days after lifting to the air liquid interface. (B) Efficient depletion of VPS35 and VPS29 was observed in day 3 cultures, but protein levels begin to recover in day 6-7 cultures, which is quantified in the graphs to the right; two-way ANOVA with Dunnett post-hoc test from three biological repeats, and error bars are SEM.

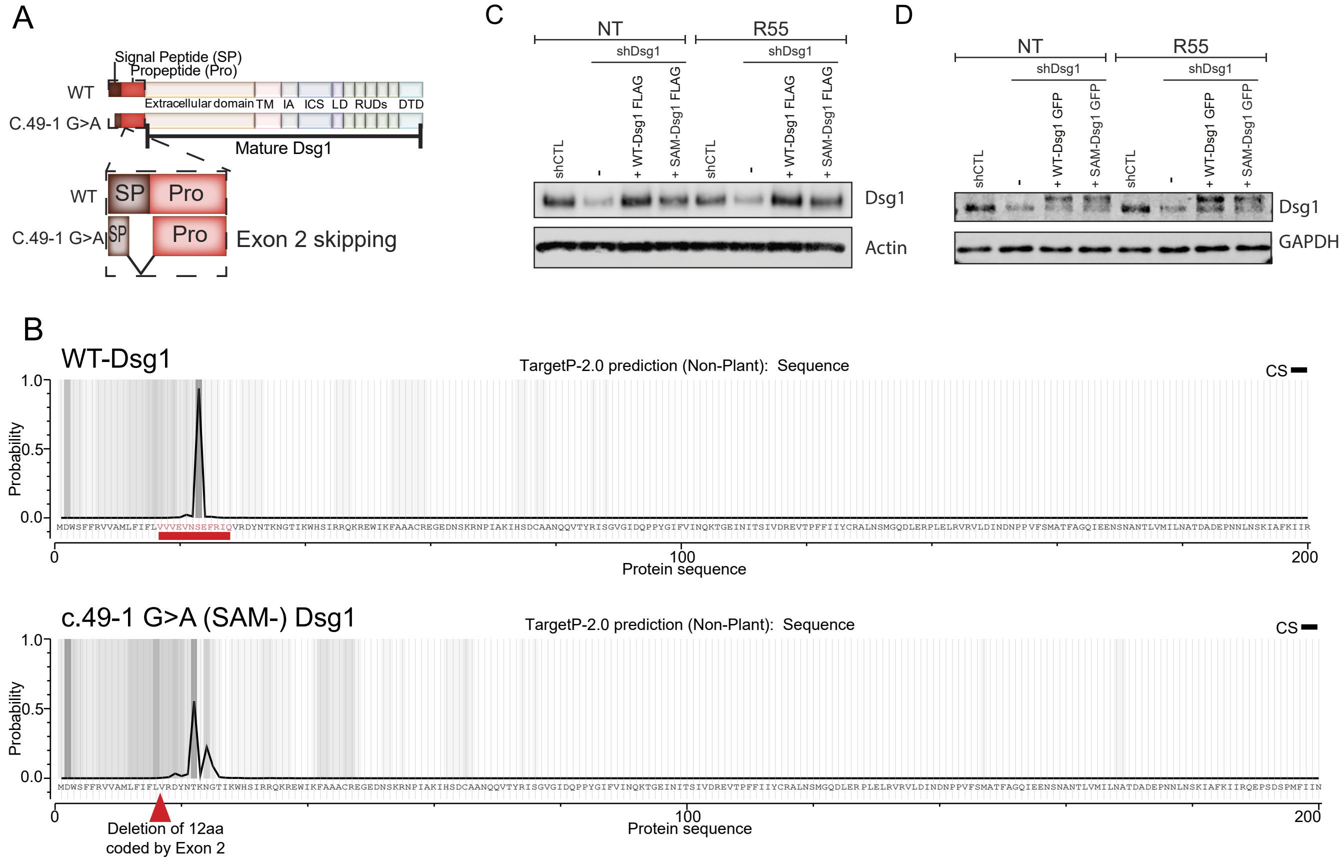


**Figure S5. Signal peptide and cleavage site prediction of WT-Dsg1 and SAM-Dsg1. R55 treatment does not affect total protein levels of Dsg1.** (A) c.49-1 G>A has been reported to result in a skipping of exon 2, which codes for amino acids located in the signal peptide sequence of Dsg1 (Samuelov et al., 2013). (B) Amino acid sequence of Dsg1with and without the deletion of exon 2 were inputted into TargetP 2.0 (http://www.cbs.dtu.dk/services/TargetP/) to test the probability of signal peptide recognition. WT-Dsg1 has a 0.981 likelihood of containing a signal peptide and a 0.9370 probability of a cleavage site (CS) at position 23-24. SAM-associated Dsg1 mutation c.49-1 G>A results in a deletion of 12 amino acids coded by Exon 2 resulting in a 0.5477 likelihood of containing a signal sequence and a 0.5527 probability that the cleavage site is between positions 22-23. The plot shows the probability and location of the cleavage site (CS), as well as the attention weight, which weights the amino acids that have higher importance to the model illustrating them with a darker shading. (C) Immunoblot of lysates expressing shCTL, shDsg1, shDsg1+WT-Dsg1-FLAG or shDsg1+SAM-Dsg1-FLAG +/-R55 and probed for Dsg1. Actin was used as a loading control. (D) Immunoblot of lysates expressing shCTL, shDsg1, shDsg1+WT-Dsg1-GFP or shDsg1+SAM-Dsg1-GFP +/-R55 and probed for Dsg1. GAPDH was used as a loading control. Whole cell lysate shown in (C) and (D) were collected in tandem with experiments shown in Figures 6 and 7, respectively.
